# Normalization of generalized transcript degradation improves accuracy in RNA-seq analysis

**DOI:** 10.1101/386938

**Authors:** Bin Xiong, Yiben Yang, Frank R. Fineis, Ji-Ping Wang

**Author notes:** Correspondence (J.-P. W.).

## Abstract

RNA-seq is a high-throughput assay to profile transcriptional activities in cells. Here we show that transcript degradation is gene-/sample-specific and presents a common and major source that may substantially bias the results in RNA-seq analysis. Most existing global normalization approaches are ineffective to correct for the degradation bias. We propose a novel pipeline named DegNorm (stands for degradation normalization) to adjust read counts for transcript degradation heterogeneity on a gene-by-gene basis while simultaneously controlling the sequencing depth. The robust and effective performance of this method is demonstrated in an extensive set of real RNA-seq data and simulated data.

## Background

RNA-seq is currently the most prevailing method for profiling transcriptional activities using high throughput sequencing technology (1). The sequencing tag count per unit of transcript length is used to measure the relative abundance of the transcript (2). Various factors exist that may affect faithful representation of transcript abundance by RNA-seq read count. Normalization is a crucial step in post-experiment data processing to ensure a fair comparison of gene expression in RNA-seq analysis (3, 4). The most commonly used approach is to normalize the read counts globally by a sample-specific scale factor for adjustment of sequencing depth. Choices of the scale factor include the total number of reads (or mean), median, trimmed mean of M-values (5), and upper quartile (3). The second type of normalization aims to remove the read count bias due to physical or chemical features of RNA sequences or uncontrollable technical aspects. The GC content is known to affect the read count in a nonlinear way (6, 7), and this effect can be sample-specific under different culture or library preparation protocols (8). Systematic bias may also arise due to technical effects such as library preparation and sequencing batches. Such systematic biases can be quantified and removed using factor analysis provided that the unwanted variation is uncorrelated with the covariates of interest (9, 10).

Another type of bias arises from cDNA fragmentation and mRNA degradation. The RNA-seq assay requires fragmenting the cDNA (reversely transcribed from mRNA) or mRNA for high throughput sequencing. Ideally, for a complete transcript, if the fragmentation is completely random, we expect to see a uniform distribution of reads along the transcript. Nevertheless the fragmentation by random priming is not really random due to primer specificity (11-13). Consequently read count per unit length of transcript may not strictly reflect the transcript abundance when comparing the expression of different genes. For the same gene, assuming the same protocol is applied to different samples, the bias attributable to fragmentation across samples should be similar. Thus fragmentation bias is less problematic in a gene-by-gene differential expression (DE) analysis. In contrast mRNA degradation can vary substantially in extent and in pattern between genes and between samples (14, 15). The degradation typically happens in the 5’ end of transcript, but it can also happen in other regions (16). Perfect control of sample degradation during the experiment is difficult, particularly when the samples are collected from field studies. More importantly, different genes may degrade at different rate (17), which makes it impossible to remove this bias by normalizing the read counts of all genes in the same sample by a same constant.

While the major impact of RNA degradation on gene expression analysis has been well recognized (17, 18) methods for correcting the degradation bias have not been fully explored in the literature. A few methods have been proposed to quantify the RNA integrity including RNA integrity numbers (RIN) (19), mRIN (20) and transcript integrity number (TIN) (13). The RIN gives a sample-specific overall RNA quality measure, but not in the gene level. The mRIN and TIN measures were both defined in the gene level by comparing the sample read distribution with reference to the hypothetical uniform distribution. In real data, due to GC-content bias, primer specificity and other complexities, the read count may substantially deviate from the uniform distribution along the transcript (7).

To reduce the degradation effect, Finotello et al. proposed to quantify exon-level expression by the maximum of its per-base counts instead of the raw read count (21). If a given exon is in a more degraded region, the local maximum may still be an under-estimate of true abundance. On the other hand, the larger variance associated with the local maximum (e.g. spikes) may result in instability in DE analysis. Based on the TIN measure, Wang et al proposed a degradation normalization method based on loess regression of read counts on TIN measure for genes within the same sample (13). However the uniform baseline assumption and the failure to compare gene-specific degradation across samples appear to be two major limitations, which may lead to extreme variability and bias in DE analysis (to be shown below).

Alternative splicing is frequently observed in higher organisms and it further complicates the gene expression estimation in RNA-seq (22). In the gene level DE analysis, we test the equivalence of relative abundance of transcripts in copy numbers between samples or conditions. If the two samples have differential exon usage, read counts need to be adjusted accordingly to better represent the transcript relative abundance in respective samples. Currently, most existing statistical packages for RNA-seq analysis (e.g., DESeq (23) and edgeR (24)) all take the raw read counts as input, while such complexities are completely ignored in practice.

In this paper, we propose a novel method to quantify transcript degradation in a generalized sense for each gene within each sample by adaptively estimating a common latent ideal coverage curve along the transcript. Using the estimated degradation index scores, we build a normalization pipeline named DegNorm to correct for degradation bias on a gene-by-gene basis while simultaneously controlling the sequencing depth. The performance of the proposed pipeline is investigated using an extensive set of real data and simulated data.

## Results

### Data sets

We consider four benchmark data sets that have been repeatedly used in the literature to assess the performance of our method. The first one was from a brain gliablastoma (GBM) cell line study of human for impact of RNA degradation (25). There were 9 samples in three groups of three, corresponding to three average RNA integrity number (RIN) = 10, 6 and 4 respectively (to be referred to as R10, R6 and R4 for simplicity). These samples were technical replicates that had undergone emulated degradation procedure. We will perform DE analysis for R10 vs R4 and R6 vs R4.

The second set contained 32 single-end samples from human peripheral blood mononuclear cells (PBMC) of four different subjects, S00, S01, S02, and S03 (17). The extracted RNA sample from each subject was kept in room temperature for 0, 12, 24, 36, 48, 60, 72 and 84 hours respectively to approximate the natural degradation process. We choose S01 as illustrating example and will perform DE analysis for 0 +12 hours vs 24 +48 hours (results for other subjects are similar).

The third set was from Sequencing Quality Control (SEQC) Consortium (26, 27) and contained two subsets, namely SEQC-AA and SEQC-AB. The SEQC-AA subset consisted of 16 technical replicates from Stratagene's universal human reference (UHR) RNA library with 2 runs for 8 lanes each. We will run DE analysis of first run vs the second run. The second subset contained two biological conditions, condition A of 5 samples from the same Stratagene's UHR RNA, and condition B of 5 samples from Ambion's human brain reference RNA. The first four replicates from both conditions were prepared by the same technician while the fifth were by Illumina. We excluded the fifth sample from both conditions because they showed dramatic difference in coverage curves compared to the rest.

The fourth data set contained 6 paired-end RNA-seq samples for embryonic zebrafish cells, three for treatment and three for control (10). There were 92 spike-in controls included in each sample while only 55 were detected present using the RUVSeq package (10) in this study.

### Nonuniformity and heterogeneity in read distribution pattern

We first define the total transcript as the concatenation of all annotated exons from the same gene. The read coverage score at a given location within the transcript is defined as the total number of reads (single-end) or DNA fragments (paired-end) that cover this position (**Supplementary Methods**). If mRNA transcripts are complete and the fragmentation is random, we expect to see a flat coverage curve in the entire transcript except the head and tail region (**Fig. 1a**). Nevertheless, in real data the read coverage curves rarely display a uniform pattern, instead dramatic and gene-specific differences are often observed across samples (**Fig. 1b-e**). The non-uniformity itself is less concerning as long as the coverage pattern is consistent across samples (**Fig. 1b**) such that the read counts can still faithfully represent the relative abundance of transcripts. In contrast, heterogeneous coverage patterns are often observed where some samples show significantly decayed read count in some regions (**Fig. 1c, d, e**), even for the spike-in controls (**Fig. 1c**). One major cause of this heterogeneity is mRNA degradation, which is clearly shown in the case of ACTN4 gene of R4 samples from the GBM data (**Fig. 1d**). Alternative splicing may also result in depleted read count in the entire region of an exon, as exemplified ST7 gene of B samples (region 1,900 - 2,900 bp) from the SEQC-AB (**Fig. 1e**). For the gene level DE analysis loss of read count due to such complexities needs to be compensated to ensure unbiased quantification of gene expression.

**Figure 1.**
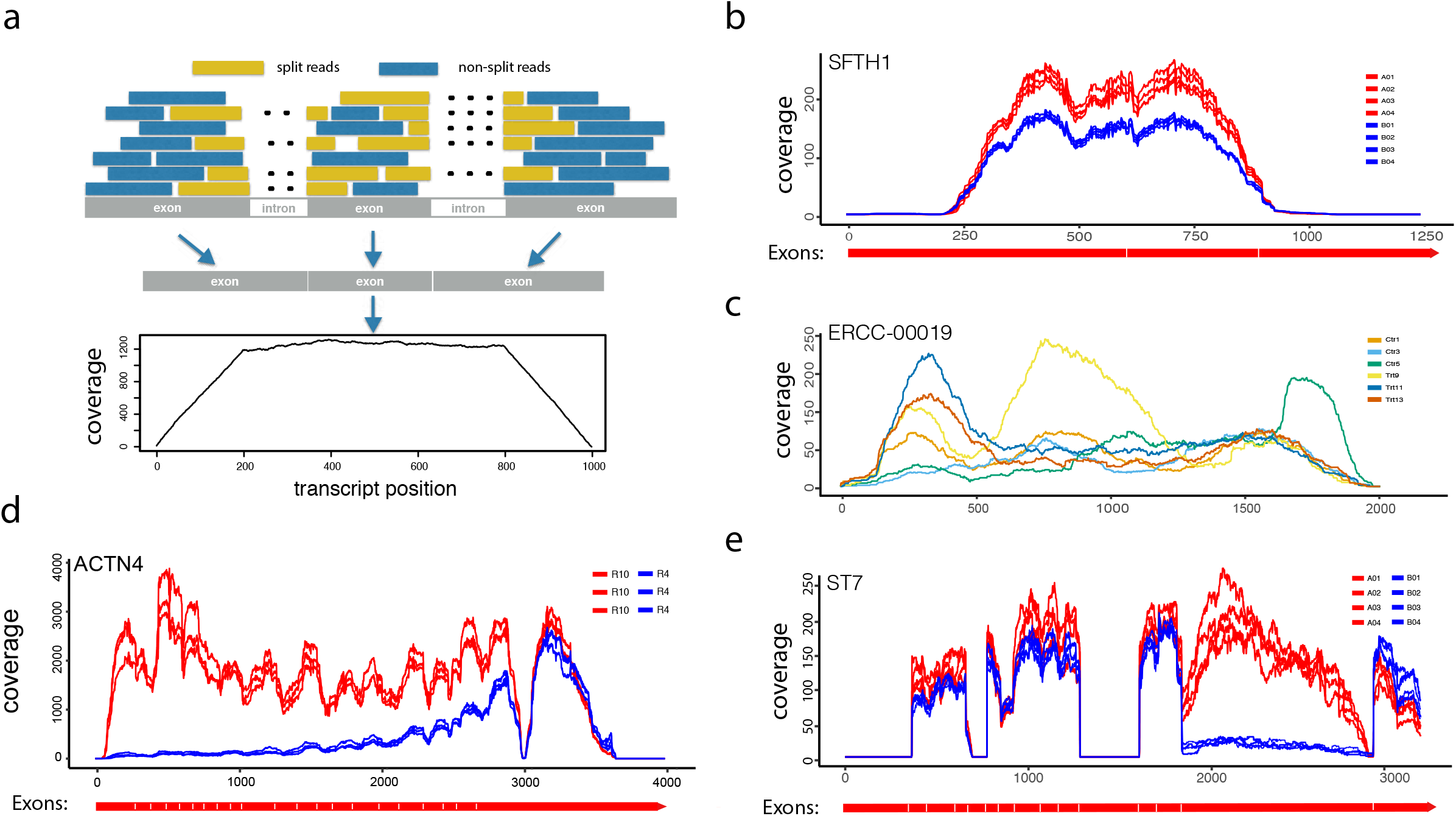
RNA-seq read coverage score shows between-sample heterogeneity in pattern along transcripts. (**a**) The read coverage score, defined as the number of reads that cover each base pair, is expected to have a trapezoidal shape along the transcript if the read start position is uniformly distributed. (**b**) An example from SEQC-AB data shows a non-uniform but consistent read coverage pattern, where the average magnitude of coverage score of each sample may faithfully represent the transcript abundance given the sequencing depth is normalized. The diagram in red under the coverage plot shows the total transcript with exon boundaries from genome annotations (same for d and e below). (**c**) An ERCC spike-in control from Zebrafish data shows non-uniform and heterogeneous coverage curve patterns between samples. (**d**) ACTN4 gene from GBM data with RIN number =10 vs RIN = 4 shows clear degraded coverage score towards to the 5’ end of the transcripts in the latter group. (**e**) An example from SEQC-AB data shows that alternative splicing likely causes sharply decayed coverage score in the entire alternatively spliced exon region.

### Generalized degradation and degradation normalization algorithm

The degradation we target to normalize is defined in a generalized sense. Any systematic decay of read count in any region of a transcript in one or more samples compared to the rest in the same study is regarded as degradation. Clearly mRNA degradation is one main cause, but alternative splicing and other factors may be the confounders that are difficult to deconvolute. To avoid confusion, in the following context we will reserve the term “mRNA degradation” for the physical degradation of mRNA sequences, and “degradation” or “transcript degradation” for the generalized degradation without specification.

We propose a degradation normalization pipeline DegNorm based on nonnegative matrix factorization over approximation (NMF-OA, See **Methods and Supplementary Methods**). We assume there is a gene-specific ideal shape of coverage curve, called an “envelope” function, identical across samples in the given study. Each envelope function is scaled by a sample- and gene-specific abundance parameter to represent the expected coverage curve for the given gene within each sample if no degradation occurs. Degradation may occur in any region of the transcript to cause negative bias in observed read counts. To illustrate this, we generated four expected coverage curves of identical shape but with different abundance levels (**Fig. 2a**), among which samples S1 and S2 are subject to degradation in the 5’ end with different patterns. Based on the expected curves, we further simulated a random realization of four complete curves with sampling error imposed (**Fig. 2b**, to be referred to as latent curves), and two degraded for sample S1 and S2 respectively (**Fig. 2c**). The NMF-OA algorithm takes the four observed coverage curves ( i.e., two non-degraded (S3 and S4) and two degraded (S1 and S2)) as input, and estimates the latent curves by minimizing the squared distance between the observed and latent, subject to the constraint that the latent curves must dominate respective observed curves at all positions (**Figs. 2d, e, Methods**). We define the degradation index (DI) score, as the fraction of area covered by the estimated latent curve, but not by the observed curve (**Figs. 2e**). It measures the proportion of missing read count due to degradation given the current sequencing depth.

**Figure 2.**
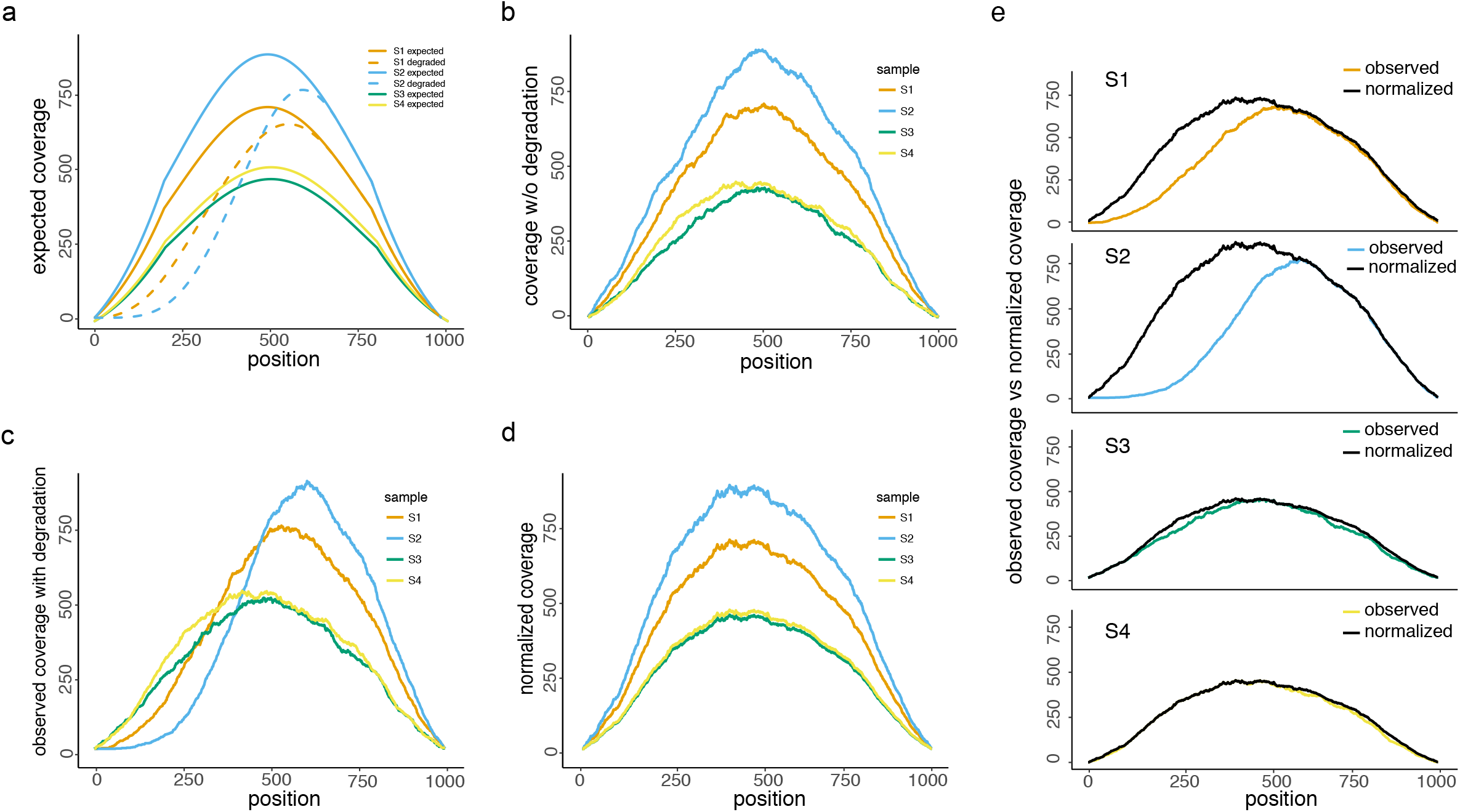
A proof-of-concept example for the proposed method for normalizing degradation patterns. (**a**) Expected or theoretical read coverage curves of one gene from 4 samples of identical shape (solid lines, without degradation), 2 of which (S1 and S2) are subject to degradation according to a rate indicated by the dashed lines in the 5’ end. (**b**) A realization of the four coverage curves without degradation randomly simulated according to the expected curves in (**a**). These curves are regarded as the latent and unobserved data. (**c**) Observed coverage curves after imposing random degradation to S1 and S2 (S3 and S4 stay intact). (**d**) Estimates of the non-degraded latent curves from the proposed algorithm solely based on the observed coverage curves in (**c**). (**e**) A sample-by-sample comparison between observed and estimated latent coverage curves.

We developed a pipeline named DegNorm that iteratively corrects for degradation bias while simultaneously normalizing sequencing depth. First, the NMF-OA algorithm is applied to sequencing-depth normalized read counts for all genes one by one to estimate the DI scores. Second, the resulting DI scores are then used to adjust the read counts by extrapolation for each gene. The adjusted read counts are used to normalize the raw data for sequencing depth. These two steps are repeated until the algorithm converges (**Methods**).

### DI score as sample quality diagnostics

The estimated DI scores provide an overview of the within-sample, between-sample and between-condition variation of degradation extent and patterns. We plotted the DI scores in three ways: a box plot of DI scores for each sample (**Figs. 3a-e**); a heatmap of the DI scores sorted in ascending order of the average scores of the first condition defined in the DE analysis (**Figs. 3f-j**); and a pairwise correlation matrix of DI scores between samples (**Supplementary Figs. 1a-e**).

**Figure 3.**
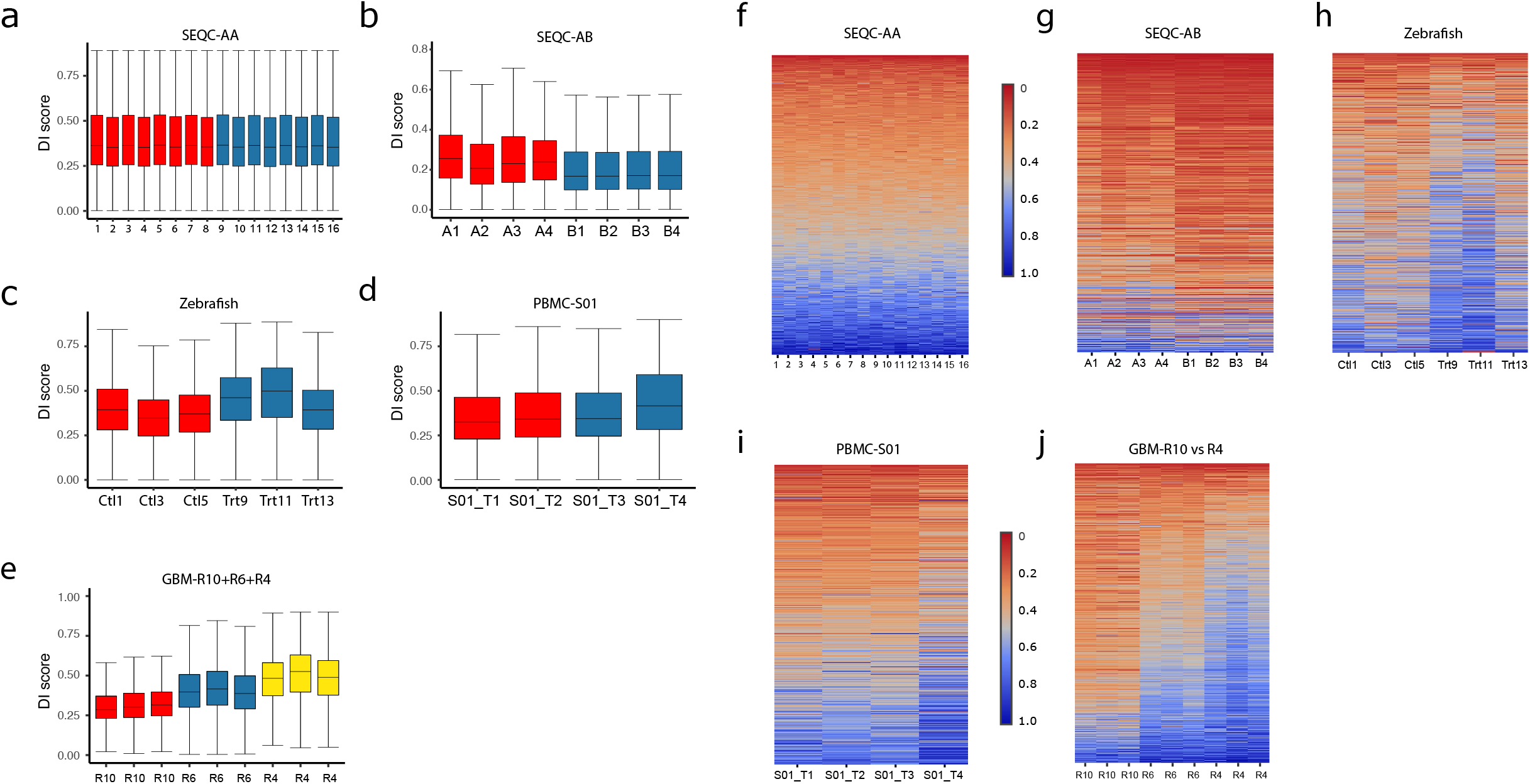
Degradation index (DI) scores show gene-/sample-/condition-specific degradation heterogeneity. (**a-f**) Box plots of DI scores presented in a between-group comparison defined for the differential expression analysis as follows. (**a**) SEQC-AA data (25,262 genes) : the eight (1-8) technical replicates from the first run vs. the eight (9-16) from the second run. (**b**) SEQC-AB data (19,061 genes): four biological replicates from A condition vs. four from condition B. (**c**) Zebrafish data (14,454 genes): three samples from the control vs. three from the treatment. (**d**) PBMC data for Subject S01 (14,051 genes): two samples exposed at room temperature for 0 and 12 hours (S01_T1 and S01_T2) vs. two for 24 and 48 hours (S01_T3 and S01_T4) respectively. (**e**) GBM data (14,298 genes): three replicates each for RIN number =10, 6 and 4 respectively. (**f-j**) For each data set presented in (**a-f**), the heatmap presents the DI scores of genes sorted in the ascending order of the average DI score of the first condition (in the GBM case, the R10 samples), where each row corresponds to the same gene across samples.

For SEQC-AA data with 16 technical replicates the median of DI scores is ~ 0.35, consistent across samples (**Fig. 3a**). While degradation pattern varies between genes, no systematic between-condition difference is observed (**Fig. 3f, Supplementary Fig. 1a**). In contrast, for the SEQC-AB data, the 4 samples from condition B have relative lower and more homogeneous degradation than A samples (**Fig. 3b**). Many genes show a condition-specific clustered pattern in DI scores (**Fig. 3g**), resulting in high within-condition correlation (**Supplementary Fig. 1b**). For the Zebrafish data, the treatment samples have slightly higher degradation than the control samples (**Fig. 3c**), while no condition-specific strong correlation is observed (**Figs. 3h, Supplementary Fig. 1c**).

The PBMC and GBM data are known to have differential mRNA degradation. The DI scores of PBMC S01 data confirm a progressive deterioration of average degradation when samples underwent degradation in the room temperature for 0, 12, 24 and 48 hours respectively (**Figs. 3d, i, Supplementary Fig. 1d**). The degradation from 24 to 48 hour was particularly accelerated than the first 24 hours. The nine GBM samples were previously classified into three groups according to the RNA Integrity Number (RIN), R = 10, 6 and 4 respectively. The DI scores show a clear ladder of escalating severity of degradation of the three groups (**Figs. 3e, j**) with two strong clusters, i.e., R10 vs. R6+R4 (**Supplementary Fig. 1e**).

In summary the DI scores from the DegNorm provide meaningful quantification of gene-level degradation between samples. The degradation pattern is gene-specific, and degradation extent may vary substantially between samples or conditions.

### DegNorm improves accuracy in gene expression analysis

We set to evaluate how the proposed DegNorm pipeline may improve DE analysis by comparing it with four other normalization methods including UQ (3), TIN (13), RUVr and RUVg (10). The RUV methods were designed to remove unwanted variation, but it is unclear whether it is effective to correct for the degradation bias. We dropped the RUVg method from the main text for its performance can be very sensitive to the choice of empirical control genes (**Supplementary Figs. 2a-h**) or the choice of factor(s) from the factor analysis in the estimation of unwanted variation (**Supplementary Figs. 2i, j**). We first examine the three data sets that originated from samples that had no true biological difference (i.e., SEQC-AA, PBMC-S01 and GBM). RNA Degradation induces bias and thus may cause extra variance or even true differential expression if some transcripts are completely degraded. Thus we investigate how different methods may reduce variance by plotting the coefficient of variation (CV) of normalized read count vs. mean read count in log scale ( the log of mean was linearly transformed to 0-1 range, **Figs. 4a-d**). In all four comparisons, the TIN method gives the largest CV than the other three methods. When RNA degradation is a major concern in GBM-R10vsR4, DegNorm substantially reduced CV compared to other methods except in the very lower end where CV was inflated due to outliers (**Fig. 4d**). The RUVr approach applies UQ normalization first, and then further removes additional variation estimated from the factor analysis. It always reduces CV over the UQ method.

**Figure 4.**
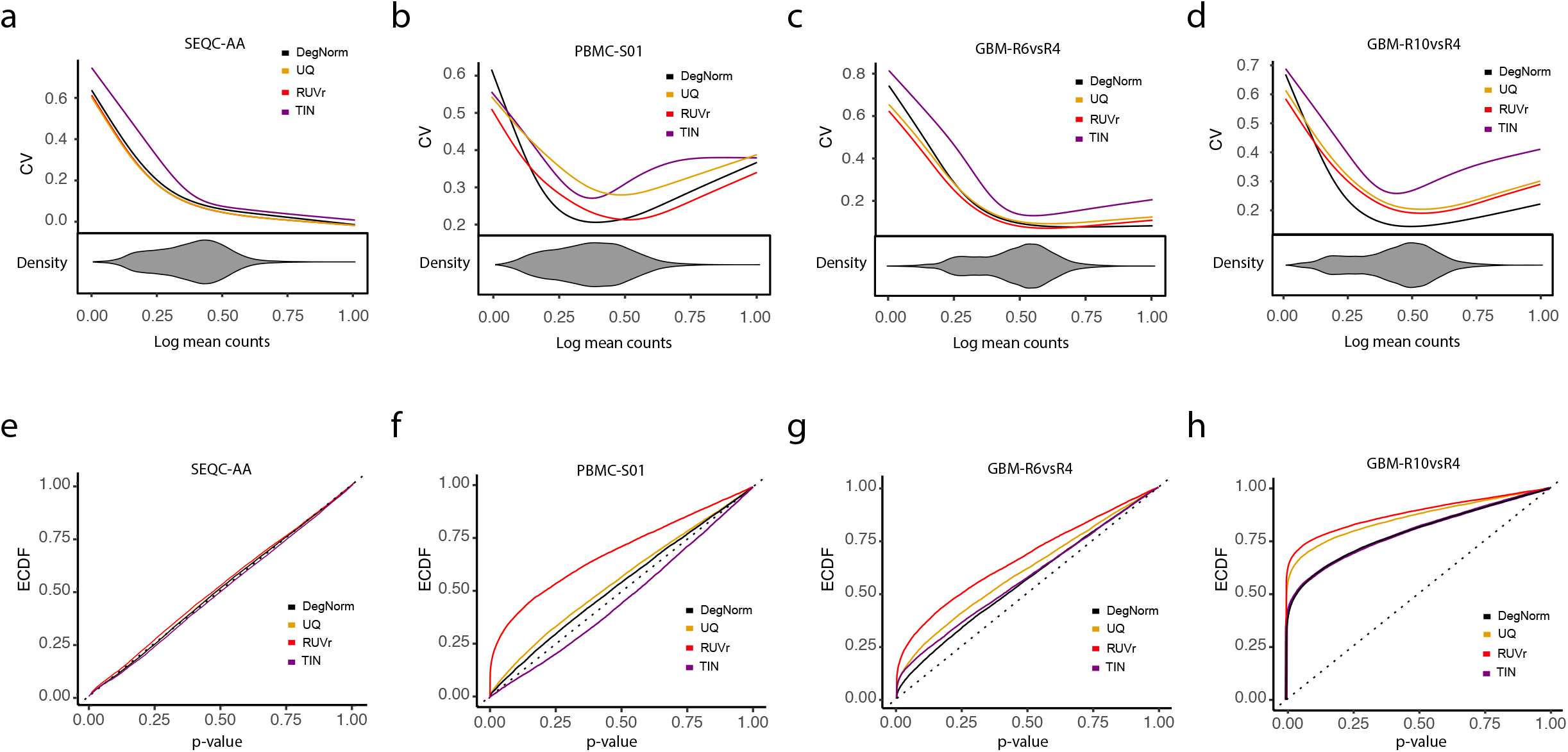
Differential expression (DE) analysis in data sets that are technical replicates or technical replicates that underwent differential degradation. Results are shown for SEQC-AA, PBMC-S01, GBM R10 + R6 and GBM R10 + R4. (**a-d**) Coefficient of variation (CV) vs. mean read count (in log scale): compared are results from proposed DegNorm pipeline and other methods including upper quartile (UQ), RUVr and TIN. The mean counts for each data were scaled to 0-1 range by a linear transformation (i.e., (*X_i_ - minj*(*X_j_*))/*max_j_* (*X_j_*) where *X_i_* is the log of the mean count for gene *i*). The CV curve was generated using the R built-in function smoothing.spline. The beanplot under the CV plot shows the density of log of mean read counts from DegNorm (The densities of read counts from other normalization methods are similar, and not shown). (**e-h**) Empirical cumulative distribution function (ECDF) of the p-value from DE analysis. The RUVr results were generated using the RUV-seq package. For TIN method, we followed Wang et al. (13) with details described in Supplementary Materials.

To further investigate how different normalization methods may improve DE analysis, we plotted the empirical cumulative distribution function (ECDF) of p-value from edgeR package. For SEQC-AA data with 16 technical replicates, all five methods gave expected ECDF close to the diagonal line (**Fig. 4e**). For PBMC-S01 and GBM-R6vsR4 comparisons with known but mild mRNA degradation variation, the ECDF from different methods were all well above the diagonal line (except TIN for PBMC data), suggesting that degradation probably have caused true differentially expressed genes (**Figs. 4f, g**). Both DegNorm and TIN methods brought the ECDF curve down towards the diagonal line compared to UQ, indicating that correction for degradation bias help reduce potential false positives. Nevertheless the TIN curve in PBMC-S01 data was well-below the diagonal line (**Fig. 4f**), which may indicate a loss of statistical power due to large variance of normalized read counts (**Fig. 4b**, to be further illustrated below). In contrast the RUVr ECDF curves in both comparisons are significantly higher than UQ (regardless that RUVr had lower CV than UQ), suggesting an ineffective correction of degradation bias or even an adverse effect to cause false positives. For the GBM-R10vsR4 comparison, ECDF curves are all far above the diagonal line, indicating a substantial rate of true positives due to drastic degradation in R4 samples (**Fig. 4h**). The TIN and DegNorm methods again appear to be more effective to correct for degradation, resulting in ECDF curves closer to the diagonal line.

The SEQC-AB and Zebrafish data present comparisons of samples with true biological differences. In both cases DegNorm produced an overall lower CV than UQ and TIN methods (**Fig. 5a,b**). The SEQC-AB data is an atypical example for containing an unusually large number of differentially expressed genes (26, 28). The DegNorm ECDF is below UQ and RUVr methods while above TIN (**Fig. 5c**). With the presence of true positives, the lower ECDF can be interpreted as a tendency to result in less false positives (good) or more false negatives (bad, less power) in the DE analysis. To investigate this, we utilized two sets of PCR verified genes, one from the original MAQC study of 843 genes (28), and the other from SEQC study of 20,801 genes (26), as the ground truth to construct receiver operating characteristic curves (ROC) (**Fig. 5d,e**). Similar as in Rissio et al ^(10)^, we defined a gene as true positive if the absolute value of log2 fold change >= 2, a true negative if <= 0.1, and undefined otherwise. For both sets, the ROC curves suggest that DegNorm method achieved better true positive rate than the UQ, RUVr and TIN while controlling the false positive rate. When we vary the threshold value from 1 to 5 to define the true positive, the area under the ROC curve (AUC) from all methods increases as expected, while AUC from DegNorm remains the largest and the gap between DegNorm and other methods enlarges (**Fig. 5f**). As the true negative set is fixed in this experiment, the true positive rate drives the change of AUC as the threshold value increases. This suggests DegNorm improves the power to identify highly differentially expressed genes than other normalization methods while controlling false positive rate. Therefore we conclude that the lower ECDF curve from DegNorm method (**Fig. 5c**) manifests a good tendency to reduce false positive rate without sacrifice of statistical power.

**Figure 5.**
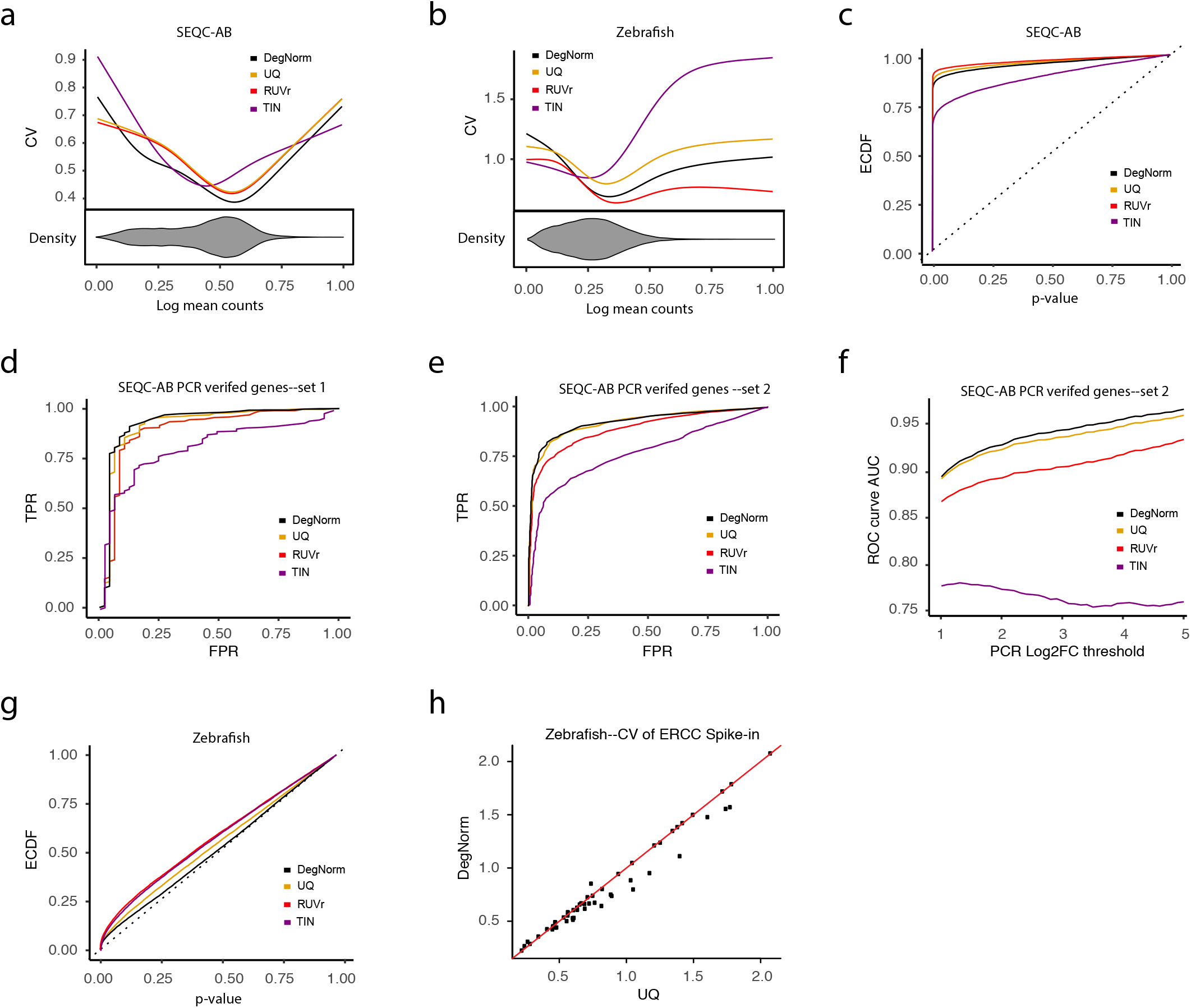
Differential expression analysis of in SEQC-AB and Zebrafish data. (**a, b**): Coefficient of variation (CV) vs. mean normalized read count (in log scale) for SEQC-AB and Zebrafish data. The beanplot under the CV plot shows the density of log of mean read counts from DegNorm. (**c**) Empirical cumulative distribution function (ECDF) of the p-value from DE analysis for SEQC-AB data. (**d**) Receiving operating characteristic curve (ROC) calculated based ~ 500 PCR verified genes (28). Genes with absolute log fold change value > 2 were defined as true positives, and absolute log fold change value < 0.1 were defined as true natives. Genes with log fold change in between were disregarded. (**e**) ROC calculated based ~ 17,304 PCR verified genes published in a separate study (26) for the same SEQC AB samples. The same threshold values of log fold change as in (**c**) were used in defining the positives and negatives. (**f**) The area under the ROC curve (AUC) statistic as a function of threshold value of absolute log fold change used to define the true positives based on the PCR data from (**e**). (**g**) Empirical cumulative distribution function (ECDF) of the p-value for Zebrafish data. (**h**) Scatter plot of CV of 55 ERCC spike-in controls in the Zebrafish data obtained using two normalization methods: DegNorm vs. UQ.

The RUVr and TIN methods both showed pronounced lower power relative to other methods, but due to different reasons (**Figs. 5d-f**). The RUVr method is guaranteed to reduce more variation based on the UQ-normalized data. Nevertheless the removed variation may contain true biological difference if it is confounded with unwanted variation, which will result in more false negatives. On the other hand, the large variance and bias incurred by TIN method appear to severely undermine the power in the DE analysis (**Figs. 4b, 5a**).

For the Zebrafish data, the ECDF from DegNorm was also the lowest among the four methods (**Fig. 5g**). As there is no verified gene set to use as ground truth to assess the overall performance, we compared the coefficient of variation of 55 ERCC spike-in controls from DegNorm and UQ methods instead (**Fig. 5h**). These spike-ins were true negatives incorporated in the treatment and control samples. About one-third of the spike-ins show a pronounced decrease of CV from DegNorm, with the rest mostly positioned close to the diagonal line. This demonstrates that normalizing degradation pattern between samples (**Figs. 3c, h**) helps remove unwanted variation due to degradation bias, and thus potentially to reduce type I error rate and improve the power as illustrated in the SEQC-AB data .

We present three examples to exemplify how DegNorm may improve the accuracy in the DE analysis. We claim a gene as positive if the local false discovery rate (lfdr) from q-value package is <= 0.05 (29-31). The MMP14 gene in the GBM-R10vsR4 comparison displayed a clear degradation in the 5’ end in R4 samples (**Fig. 6a**), and was tested positive using UQ method (p-value = 0.006, lfdr = 0.030). DegNorm compensated the degraded portion of R4 samples and returned negative test result (p-value = 0.031, lfdr = 0.128, **Fig. 6b**). The second example is the TPMT gene from SEQC-AB comparison, PCR verified positive with log fold change= 18.23. It was test-negative under UQ (p-value = 0.139, lfdr = 0.368) (**Fig. 6c**) as opposed to test-positive under DegNorm with degradation correction in the 5’ end of A samples (p-value = 1.88e-5, lfdr = 1.24e-4, **Fig. 6d**). The third example is the NDUF3 gene from the SEQC-AB data, showing nearly depleted coverage in the entire third exon region from ~223-1,327 nt for B samples likely due to alternative splicing. It was tested positive under UQ (p-value=1.94e-37, lfdr =3.39e-09, **Fig. 6e**). DegNorm returned a negative result (p-value = 0.808, lfdr = 0.942, **Fig. 6f**), consistent with negative PCR verification (log fold change = −0.114).

**Figure 6.**
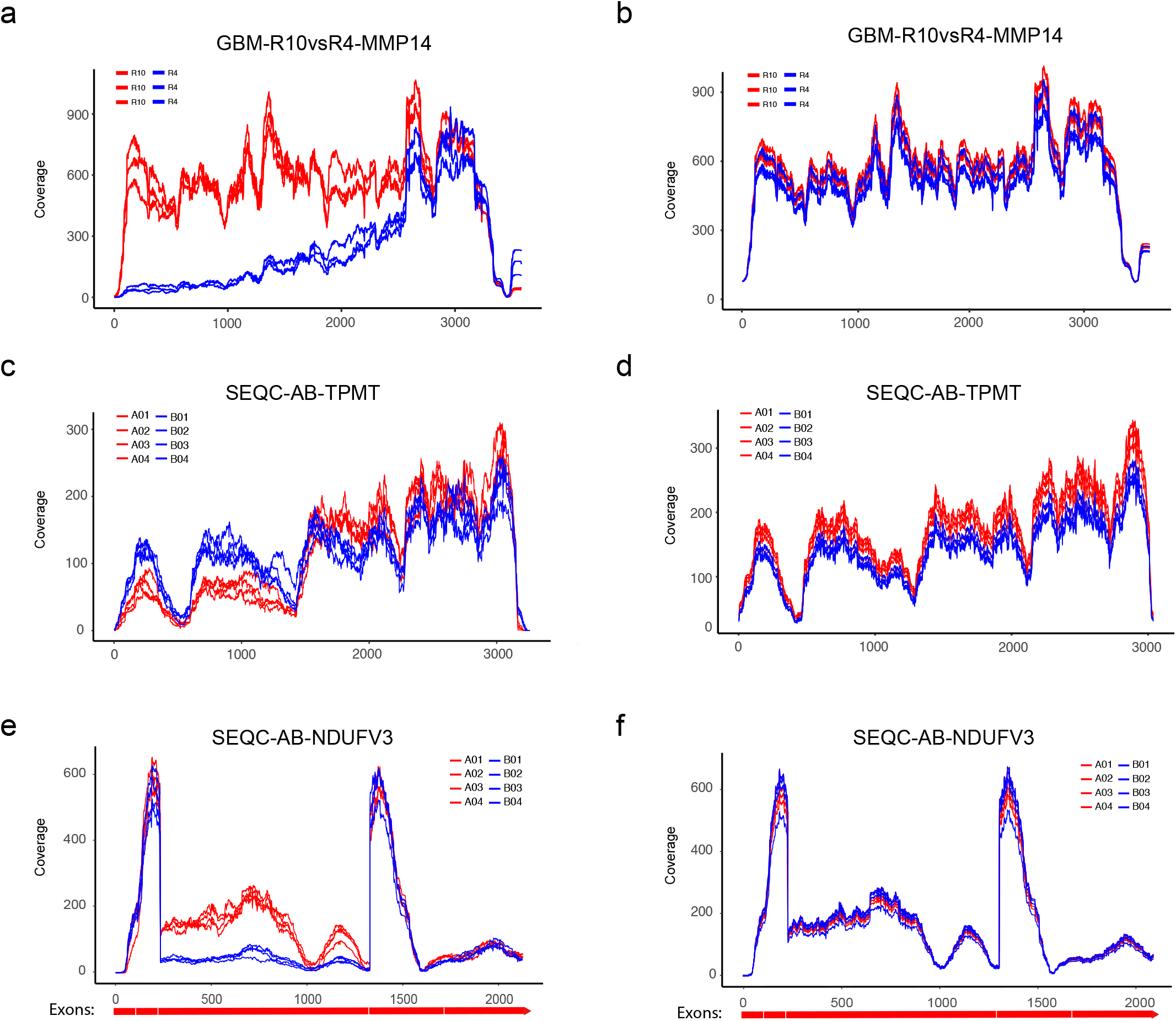
Examples illustrating that DegNorm improves accuracy in RNA-seq DE analysis. (**a, b**) Coverage curves under UQ and DegNorm normalization methods side by side for gene MMP14 in GBM R10vsR4 comparison. (**c, d**) and (**e, f**) Same coverage plots for TPMT and NDUFV3 genes respectively in the SEQC-AB comparison. Both MMP14 and NDUFV3 changed from positive to negative after degradation normalization, whereas TPMT changed from negative to positive in the DE analysis using edgeR. In (**e, f**), annotated exon boundaries are plotted to show possible alternative splicing in the exon from 223-1327 nt.

We further compared other normalization methods including trimmed mean of M-values (TMM) (5), relative log expression (RLE) (23) and total read count (TC) (4) in the comparison. The resulting ECDF plots were very similar to the UQ method in all data sets we analyzed in this paper (**Supplementary Figs. 3 a-f**).

### Simulation study

To systematically investigate the performance of DegNorm, we conducted a simulation study in a two-condition comparison: 4 control (A) vs. 4 treatment (B) samples in four different degradation settings. Each sample of 20,000 genes was simulated with a random sequencing depth of 40-60 million reads, 5% of which were chosen to be up-regulated and 5% for down-regulated in expression. In the first setting all genes have no degradation whereas in the rest three settings, 80% genes were randomly selected for degradation. In the second setting, for a gene selected for degradation, 3, 4 or 5 samples out of the 8 were randomly chosen for degradation whereas in the third, either all four control samples or four treatment samples were randomly chosen for degradation. In both second and third settings, the degradation extent for each degraded gene was random but following the same distribution. In the fourth setting, for each gene selected to degrade, two control samples were randomly selected for degradation with the same expected severity, while all treatment samples underwent degradation with sample-specific systematic difference in expected severity. The simulation details are described in **Methods** and **Supplementary Materials**.

Simulation I presents a case that samples have no degradation bias or any other bias but only between-sample variation in sequencing depth. DegNorm is data-driven and always returns nonnegative DI scores by design. The estimated DI scores have a median ~ 0.11 in all samples (**Fig. 7a**), showing no between-condition heterogeneity (**Fig. 7b**). To investigate how the positive bias in DI scores may impact DE analysis, we first plotted the normalized vs. latent read count from different methods (**Fig. 7c-f**). Unsurprisingly the UQ method perfectly normalizes sequencing depth (**Fig. 7c**), while all other three methods caused bias or extra variance to different extents (**Fig. 7d-f**). The positive bias from DegNorm is more pronounced when read counts are low such that read coverage curve cannot be well estimated (**Fig. 7e**). As for any gene, all samples are subject to this over-estimation bias, this bias may partially cancel off in DE analysis. As a result the ECDF and ROC curves of DegNorm almost perfectly overlap with the latent or UQ or RUVr curve (**Fig. 7g-h**), suggesting little adverse effect was caused by the positive bias in DE analysis. In contrast the TIN method clearly showed inferior performance in DE analysis due to excess of variance and large bias (**Fig. 7d**).

**Figure 7.**
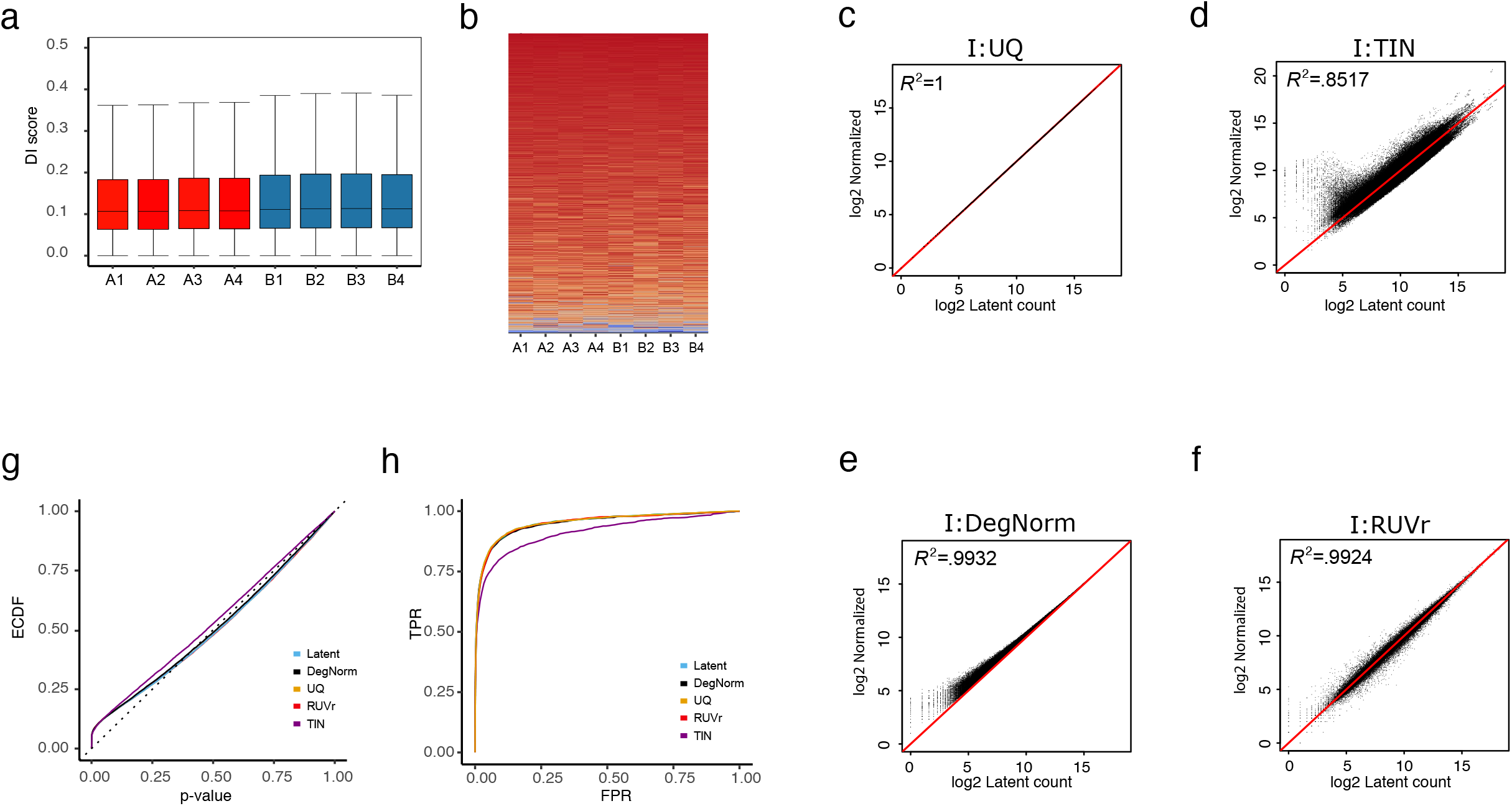
Results for Simulation I. (**a**) Box plot of estimated DI scores where condition A and B stand for control and treatment samples respectively. (**b**) Heatmap of DI scores of genes sorted in the ascending order of the average DI score of condition A (control), showing no between-condition clustered pattern. (**c-f**) Scatter plot of normalized read count vs. latent read count in log2 scale with diagonal line imposed for Simulation I. Compared are results from proposed DegNorm pipeline and other methods including upper quartile (UQ), RUVr and TIN. The mean counts for each data were scaled to 0-1 range by a linear transformation (i.e., (*X_i_ - min_j_(X_j_)*)/*max_j_ (X_j_*) where *X_i_* is the log of the mean count for gene *i*). (**g, h**) ECDF plots and ROC curves for simulation I under different normalization methods. All curves except TIN were overlapping with each other, suggesting close performance in DE analysis.

When degradation is an issue in Simulations II-IV, the estimated DI scores provide informative characterization of overall degradation severity of each sample (**Supplementary Figs. 4a-f**) as well as the similarity of gene-specific degradation pattern between samples (**Supplementary Figs. 4g-l**). DegNorm demonstrates consistently better correction of degradation bias than the UQ, TIN and RUVr methods, evidenced by higher regression coefficient of determination (*R*^2^), and a nearly symmetric distribution of data points around the diagonal line in the scatter plot of normalized vs. latent read counts (**Figs. 8a-d, Supplementary Figs. 5a-h**). The UQ method does not correct for any degradation bias but normalizing sequencing depth (**Fig. 8a, Supplementary Figs. 5a, e**). Similar extreme bias and variance issues are also observed in the TIN method in Simulations II-IV (**Fig. 8b, Supplementary Figs. 5b, f**).

**Figure 8.**
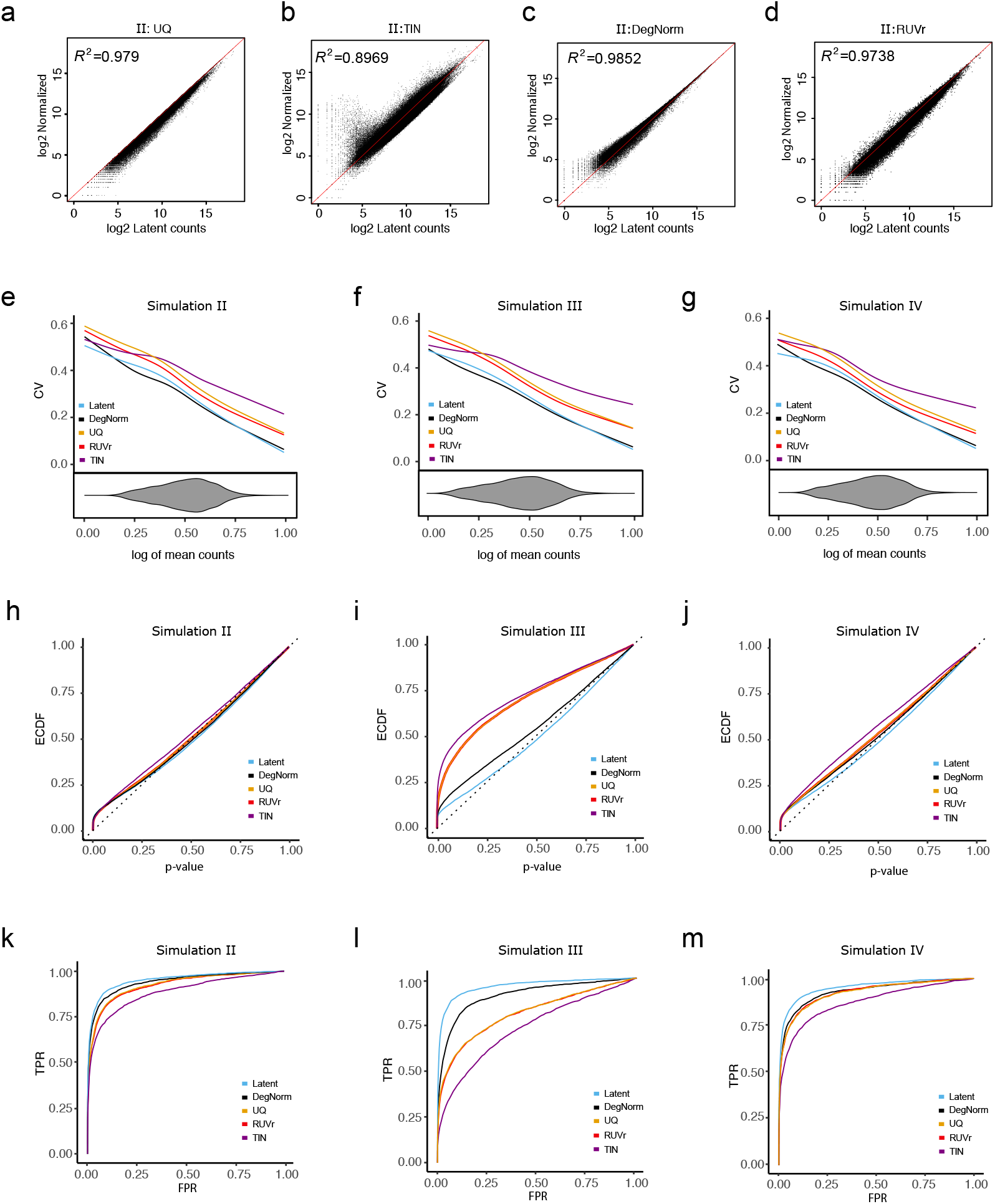
Comparison of different normalization methods in Simulation II, III and IV (**a-c**) Scatter plot of normalized read count vs. latent read count in log2 scale with diagonal line imposed from DegNorm, UQ, RUVg, RUVr and TIN methods for Simulation II. (**d-h**) Coefficient of variation (CV) vs. normalized mean read count (in log scale) for 18,000 true negative genes (out of a total of 20,000 genes). The beanplot under the CV plot shows the density of log of mean read counts from DegNorm (The densities from other normalization methods are similar, and not shown). (**i-j**) Empirical CDF of p-value from edgeR DE analysis . The latent curve corresponds to the results using the true read count before the degradation was imposed. (**k-m**) Receiver operating characteristic curves of DE analysis.

To gain more insights into the CV plots, we singled out the 18,000 true null genes from Simulations II-IV and plotted the CV vs. mean of normalized count (**Figs. 8e-g**). For the true nulls, without the confounding effect of biological difference, the CV tends to be inflated due to degradation-caused loss of read count. Thus CV is an informative measure to assess the effectiveness of degradation normalization. Indeed, the UQ and RUVr curves are both well above the latent in all simulations, suggesting degradation bias was not or inadequately corrected. Similar to observed in the real data, the TIN CV curve dominates other methods in each setting, echoing the excess of bias and variance observed in the scatter plot (**Fig. 8b, Supplementary Figs. 5b, f**). In contrast the DegNorm curves were the closest to the respective latent curves but with a slight under-estimation of CV in the lower half.

DegNorm improves the accuracy in DE analyses. In both ECDF (**Figs. 8h-j**) and ROC plots (**Figs. 8k-m**), DegNorm curve was the closest to the latent curve, showing improvement over other normalization methods to different extents. In Setting II and III, where degradation was randomly chosen among samples (II) or conditions (III), the UQ and RUVr methods were both ineffective to correct this non-systematic but gene-specific bias (**Figs. 8k, l**). In particular, in Simulation III as degradation was applied to one condition of random choice for a given gene, the degradation bias was completely confounded with covariate of interest and cannot be removed by RUVr method. Consequently many false positives tend to be called due to degradation bias (**Figs. 8l**), resulting in much worse ROC curve than the DegNorm. In contrast, the treatment samples in setting IV had systematic difference in average degradation (**Supplementary Fig. 4c**), UQ and RUVr both performed reasonably well (**Fig. 8m**). In all four degradation settings, the TIN method showed a clear undermined power or greatly inflated false positive rate.

## Discussion and conclusions

In this paper we showed that degradation pattern and severity are gene-specific, and thus commonly used global normalization methods that impose a constant adjustment to all genes within the same sample are ineffective to correct for this bias. The RUVr approach is guaranteed to reduce variation, while they failed to show pronounced improvement over the UQ method in the DE analysis in all data sets considered in this study (even got worse in PBMC and GBM data). One complexity is that the true biological difference is often confounded with degradation (e.g., SEQC-AB data) and other unwanted variation. The factor analysis cannot well separate the unwanted from the wanted variation and may even remove the true biological difference of interest. This confounding issue was illustrated in the SEQC-AB data by the high within-condition and low between-condition correlation of DI scores (**Supplementary Fig. 1b**). In particular the RUVg method was sensitive to the selection of empirical control genes or the factor(s) used to estimate the unwanted variation (**Supplementary Fig. 2a-j**). More objective criteria in this regard need to be developed for the RUVg method.

Although motivated by mRNA degradation, we defined the degradation in this paper in a generalized and relative sense. The quantified DI scores may reflect confounding effects from mRNA degradation, alternative splicing and other factors. Risso et al (10) showed that in Zebrafish and SEQC-AB data, samples were clustered due to difference of sample preparation, experiment protocol, sequencing run batches and flow cell etc. Such factors could impact the RNA samples by changing the read coverage curves. Thus normalizing heterogeneity in coverage curves may help remove bias due to such factors. Indeed in all four benchmark data sets considered in this paper, DegNorm performed consistently better compared to other methods regardless whether mRNA degradation was a known concerning issue. Unlike mRIN and TIN measures where degradation was defined as deviation from hypothesized uniform coverage curve, the DI score from DegNorm is defined with reference to an adaptively estimated latent coverage curve that minimizes the distance to the observed coverage curves. If a gene has 50% degradation but having consistent coverage curves across samples, the estimated DI scores will all be nearly 0. In this case, the read count in each sample can still accurately reflect the relative abundance between samples, and degradation correction is unnecessary. From this perspective, the DI score is defined in a relative sense. Normalizing the read count for degradation bias using DI scores is hoped to minimize the extrapolation needed, thus avoids excess of variance. The advantage of this strategy was exemplified in all real and simulated data sets in contrast to the TIN method.

There are a few limitations of the DegNorm method. First, DegNorm only corrects for bias that causes heterogeneous read coverage patterns between samples. Second, the core of DegNorm is a matrix factorization over approximation algorithm aiming to correct for the degradation bias that commonly exists in RNA samples, even for the high quality SEQC data. DegNorm tends to result in a prinounced over-estimation bias in DI scores for genes with low read counts regardless whether degradation is present (Simulations II-IV) or absent (Simulation I). We showed in Simulation I that this bias is not a big concern in DE analysis as it tends to be homogeneous across all samples for any non-degraded gene. Third, DegNorm is computing intensive due to calculation of read coverage curves and repeated singular-value-decomposition of large matrices. We are currently working on a software tool to automate the entire pipeline.

In summary we conclude DegNorm provides a pipeline for informative quantification of gene-/sample-specific transcript degradation pattern and for effective correction of degradation bias in RNA-seq. It can be practiced as a general normalization method to improve the accuracy in the gene expression differentiation analysis.

## Methods

### DegNorm algorithm

Suppose we have *p* samples and *n* genes, *X_ij_* is the read count for gene *i* in sample *j*. For simplicity of notation, we first illustrate the proposed method by focusing on one gene. Let *f_ij_*(*x*), *x* = 1,…, *L_i_*; *i* = 1,…, *n*; *j* = 1,…, *p*, be the read coverage score for transcript *i* of length *L_i_* from sample *j*. When different isoforms are present, the *L_i_* positions represent the assembly of all expressed exons in the sequential order. We assume there is an *envelope* function *e_i_*(*x*) that defines the ideal shape of read coverage curve for gene *i* if no degradation exists. The actual ideal coverage curve for the given gene in the *j*th sample is *k_ij_e_i_*(*x*), where *k_ij_* denotes the confounded effect of sequencing depth and relative abundance of gene *i* in sample *j*.

Degradation causes *f_ij_*(*x*) to deviate downward from *k_ij_e_i_*(*x*) in degraded region(s). Clearly sampling error can cause random fluctuation of *f_ij_*(*x*) from the ideal curve *k_ij_e_i_*(*x*) (**Figs. 2a, b**). We assume the random error is ignorable compared to the major bias arising from degradation. Thus we require *k_ij_e_i_*(*x*) ≥ *f_ij_*(*x*) for all *j* and *x*. The difference between *k_ij_e_i_*(*x*) and *f_ij_*(*x*) provides an estimate of degraded portion of read count. We propose a method that allows to estimate *k_ij_* and *e_i_*(*x*) to quantify the degradation extent of each gene within each sample while simultaneously controlling the sequencing depth.

### Estimating degradation via non-negative matrix over-approximation

Let **f***_ij_* = (*f_ij_*(1),…, *f_ij_*(*L_i_*))^*T*^, *j* = 1, …,*p* and **F***_i_* =(**f***_i1_*,…, **f***_ip_*)^*T*^. Let **K***_i_* = (*k_i1_*, …,*k_ip_*)^*T*^, **E***_i_* = (*e_i_*(1),…, *e_i_*(*L_i_*))^*T*^. We propose to estimate **K*_i_*** and **E*_i_*** by minimizing the following quadratic loss function subject to some constraint:

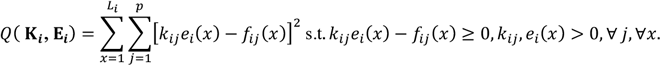

We can configure this problem into a non-negative matrix factorization problem (32, 33) as follows:

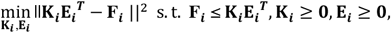

where ∥-∥^2^ stands for the element-wise quadratic norm (i.e., sum of squared elements), and ≤ and ≥ for element-wise logical comparison. We call this a rank-one non-negative matrix factorization over-approximation (NMF-OA) problem as **K*_i_*** and **E*_i_*** have rank 1 and 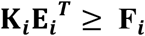. The proposed iterative algorithm is described in details in the **Supplementary Methods**.

### Refinement of NMF-OA algorithm

The NMF-OA optimization algorithm provides an approximate solution to this problem. However in the RNA-seq data, the performance of the solution can be affected by two confounding factors, degradation extent and sequencing depth. In our quadratic objective function *Q*(**K*_j_***, **E*_j_***), an **f***_ij_* of larger magnitude tends to have more influence on the estimation of envelope function. A dominant scale in **f***_ij_* may force the algorithm to fit an envelope function that resembles **f***_ij_* to minimize the loss. Thus a good scale normalizing factor for sequencing depth is important to yield a good estimate of the envelope function *e_i_*(*x*). With gene-specific and sample-specific degradation, the total number of reads may not provide a reliable measure of sequencing depth.

Second, given **F*_i_*** that is appropriately normalized for sequencing depth, the scale factor **K*_i_*** should reflect the relative abundance of the gene in the non-degraded region of each sample (to be referred as the baseline region below) (**Fig. 2a**). The non-degraded region must preserve similar shape in the coverage curve in different samples. If one can first estimate **K*_i_*** from the identified baseline region, then NMF-OA algorithm will lead to better estimate of the envelope function **E*_i_***, particularly in the situation when the degradation extent is severe as the GBM data. We account for these two considerations by proposing an iterative degradation normalization pipeline (DegNorm) as follows:

1. Sequencing depth adjustment: Given the current estimate of (**K*_i_***, **E*_i_***) for *i* = 1,…,*n*, define the degradation index (DI) score as

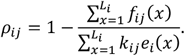 Graphically *ρ_ij_* stands for the area in fraction under the curve *k_ij_e_i_*(*x*) but above *f_ij_*(*x*) (**Fig. 2e**). The DI score is used to calculated an adjusted read count by extrapolation

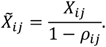 Sequencing depth scaling factor can be calculated using the degradation corrected total number of read count

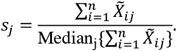 In this paper, results presented were all based on this scale normalization. Alternatively we can use other normalization methods like TMM or UQ to calculate the normalizing constant *s_j_* based on the degradation-adjusted read count 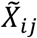 if presence of extreme outliers is a concern.
2. Degradation estimation: Given the estimated sequencing depth *s_j_* from step 1, adjust the coverage curves as follows:

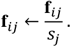

- Let **F_*i*_** =(**f**_*i*1_, …, **f**_*ip*_)^*T*^. Run NMF-OA for each gene on updated coverage curves **F*_i_***, and obtain estimate of (**K*_i_***, **E*_i_***).
- Divide each gene into 20 bins. Define the residual matrix as 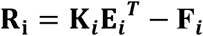 . We identify a subset of bins on which **F*_i_*** preserves the most similar shape as the envelope function **E*_i_*** across all samples by progressively dropping bins that have the largest sum of squares of normalized residuals (**R*_i_*/F*_i_***) and repeatedly applying NMF-OA to remaining bins. This step stops if the maximum DI score obtained from the remaining bins is <=0.1 or 70% bins have been dropped (see details in **Supplementary Materials**). The remaining transcript regions on selected bins are regarded as the baseline. Denote the read coverage curve on baseline region as 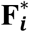.
- Run NMF-OA on 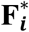. The resulting 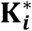 is an refined estimate of **K*_i_***, given which the envelope function can be obtained as

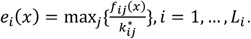
3. Steps 1 and 2 are repeated until the algorithm converges.

#### Simulations

We first simulated the latent read count for each gene within each sample. For a given gene without DE, the latent counts (without degradation) in both control and treatment samples were randomly simulated from the negative binomial distribution with the same mean and same dispersion parameters that were randomly chosen from the fitted values of SEQC-AB data. For a gene with DE, the latent counts for the control samples were first simulated as above, and those for the treatment samples were simulated from the negative binomial with mean parameter increased/decreased by a factor of (1.5 + γ) for up- or down-regulated genes respectively, where γ was simulated randomly from an exponential distribution with mean =1. In the second step, we simulated the degradation given the latent read count (Simulations II-IV). For a given gene, we first chose a Gaussian mixture distribution that covers the entire range of total transcript to model the read start position distribution. For simplicity, we only consider 5’ end degradation. Each read from a gene that was pre-selected for degradation degraded (be disregarded) with probability that depended on the start position of the read, defined by a cumulative distribution function (CDF) of a lognormal distribution. The degradation pattern and extent can be tuned by varying the parameters in the lognormal distribution (see more details in **Supplementary Methods**).

## Competing interests

The authors declare that they have no competing interests.

## Availability of data and materials

All the raw data and processed data can be accessed at http://bioinfo.stats.northwestem.edu/~jzwang/DegNormData.zip

---

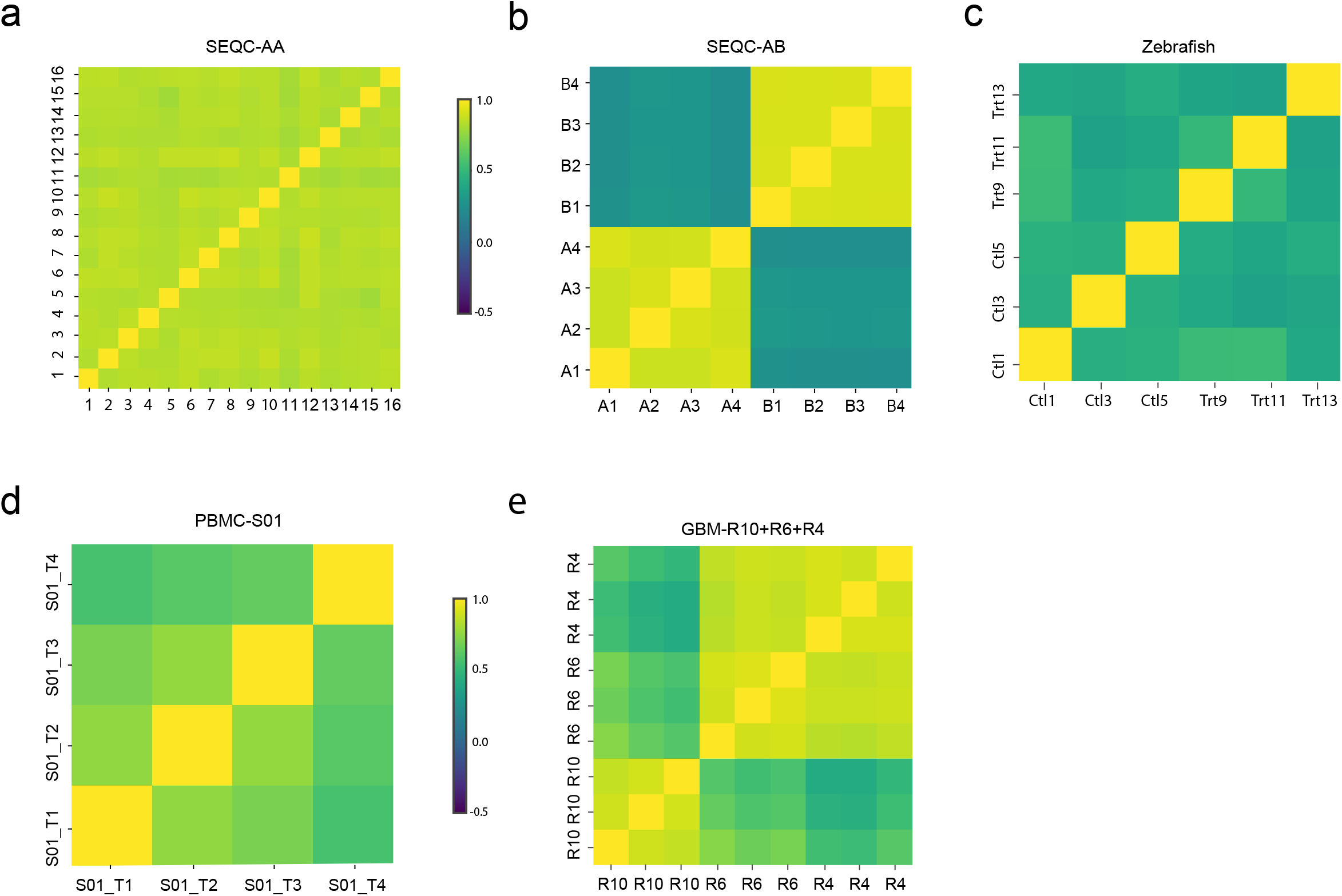

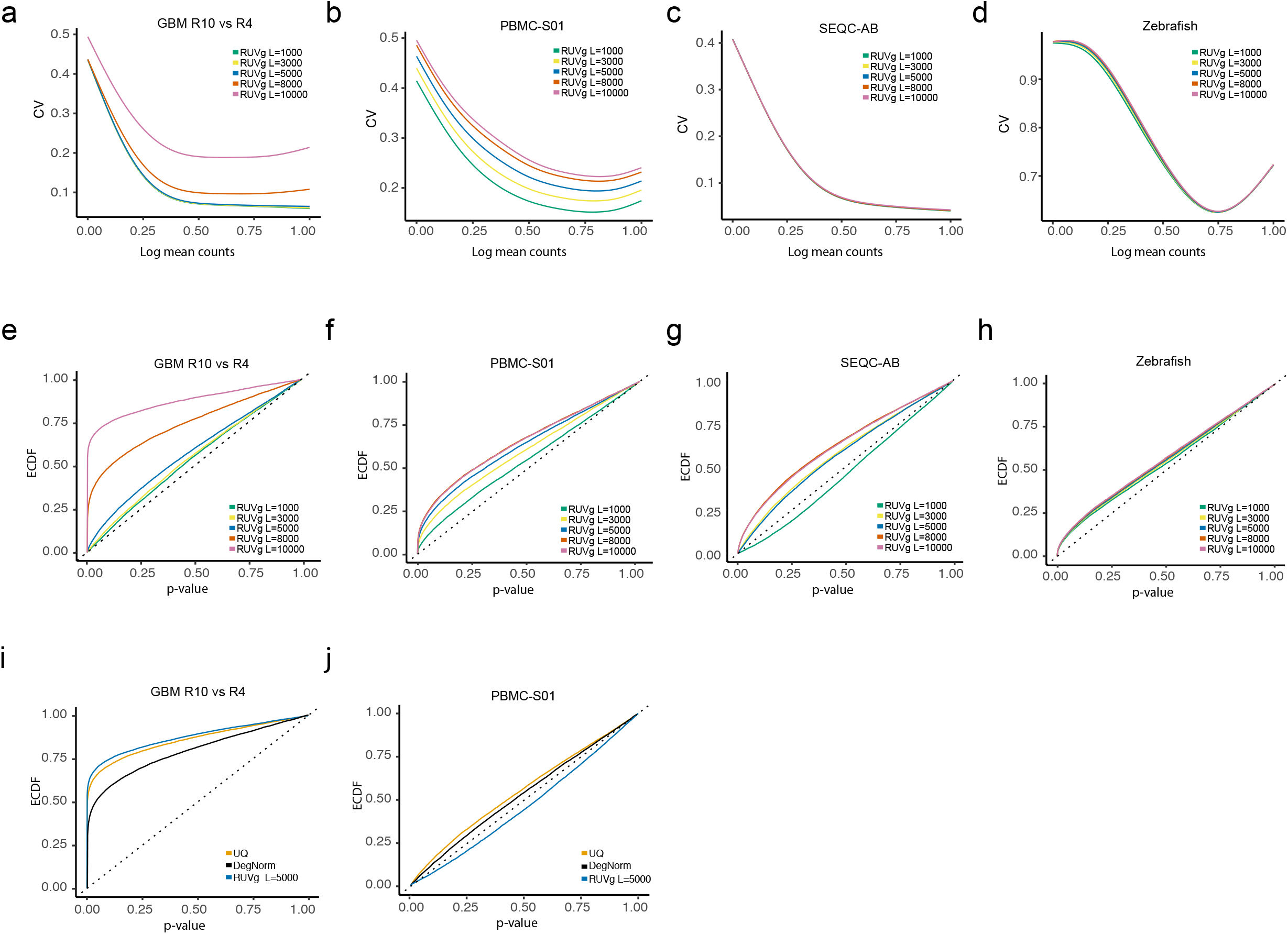

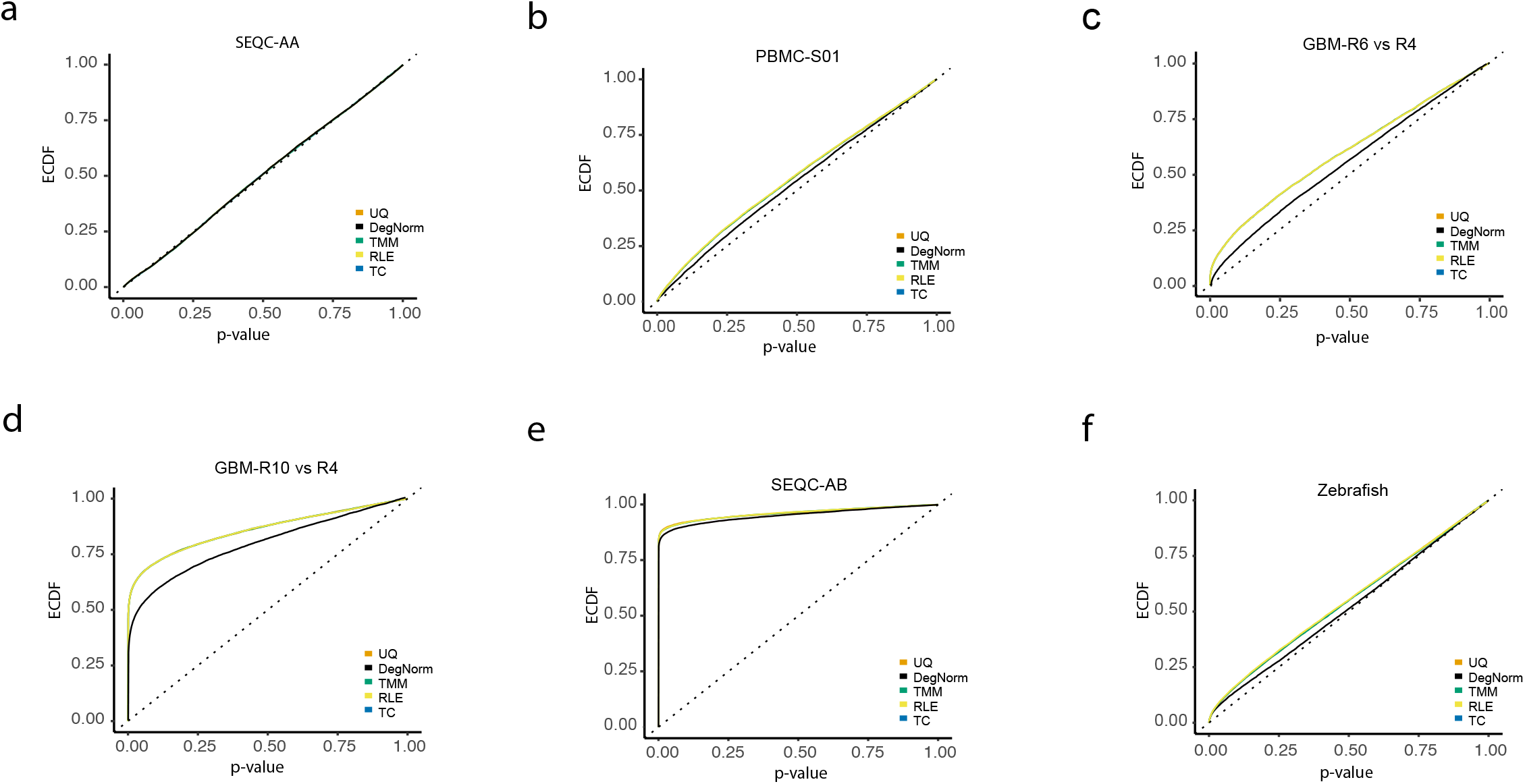

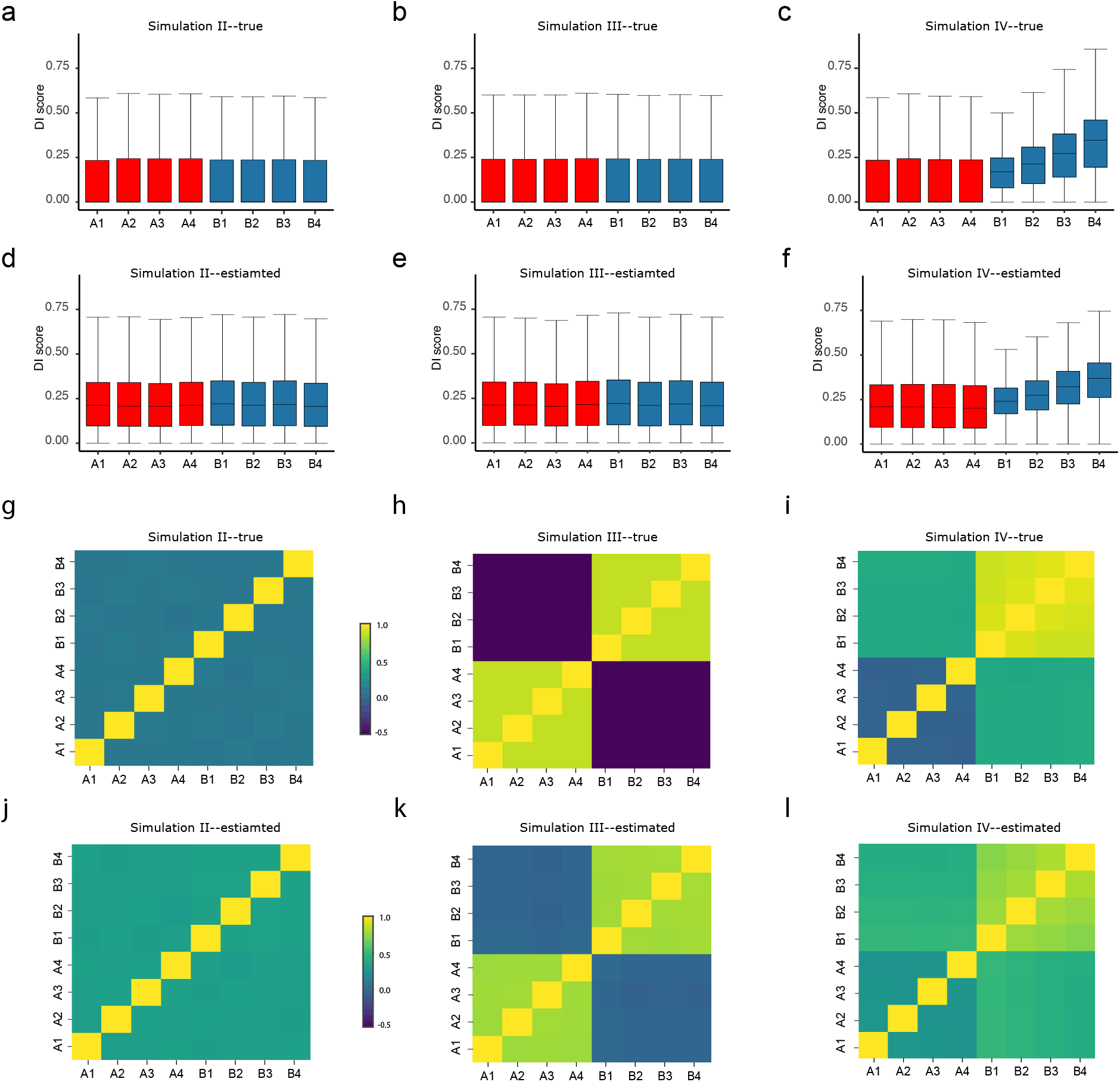

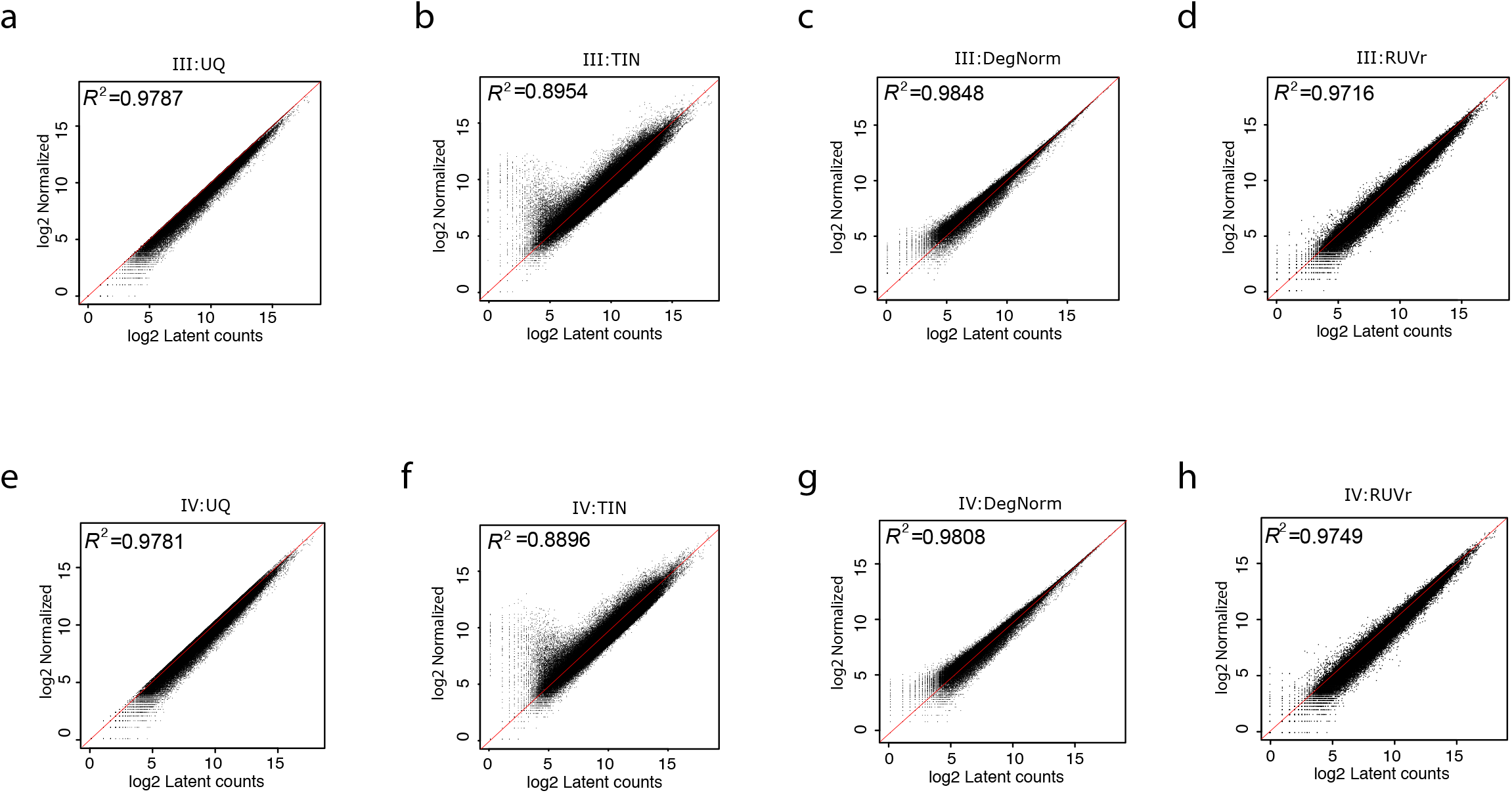

